# Optimizing Accuracy and Efficiency in Analyzing Non-UMI Liquid Biopsy Datasets Using the Sentieon ctDNA Pipeline

**DOI:** 10.1101/2024.01.24.577136

**Authors:** Li Niu, Jinnan Hu, Chuan Chen, Cai Jiang, Haodong Chen, Gongcheng Tang, Ying Liu, Yi Liu

**Affiliations:** CheerLand Clinical Laboratory Co., Ltd, Shenzhen, China; Sentieon Inc, San Jose, CA, USA; Shenzhen Cheerland Biotechnology Co., Ltd, Shenzhen, China; Nanodigmbio (Nanjing) Biotechnology Co., Ltd, Nanjing, Jiangsu, China

## Abstract

Sequencing clinical liquid biopsy, especially circulating tumor DNA (ctDNA), provides a valuable method for identifying low allele frequency tumor variants, opening novel clinical applications, particularly in treatment selection for late-stage cancer patients. Despite advancements, challenges in assay development persist, primarily due to limited sample volumes and insufficiency of reads supporting low allele frequency variants. The allele frequencies of clinically significant variants often hover close to the threshold of errors introduced by PCR and sequencing processes. Therefore, more sophisticated analysis methods are crucial to further reduce base error rates, enabling accurate discrimination between background errors and genuine somatic variants. While several ctDNA analysis pipelines have been published and adopted, there is room for improvement in terms of accuracy and run efficiency.

In this study, we introduce Sentieon’s innovative consensus-based ctDNA pipeline - a rapid and precise solution for calling small somatic variants from non-UMI ctDNA sequencing data. The pipeline comprises four core modules: alignment, consensus generation, variant calling, and variant filtering. Through benchmarking with in-vitro and real clinical datasets, we observed that the Sentieon ctDNA pipeline exhibits higher accuracy compared to alternative methods.

## Background

Tumor cells release fragments of DNA into the circulatory system during apoptosis or necrosis, known as circulating tumor DNA (ctDNA)^1^. These ctDNA fragments retain somatic variants from the original tumor tissue, allowing inference of tumor stages based on their existence and fraction^2,3^. Consequently, ctDNA detection and monitoring serve as a natural target biomarker for various clinical oncology assays, including treatment selection, minimum residual disease (MRD) monitoring, and early cancer detection^4^. Assessing somatic variants from ctDNA and other liquid biopsies offers advantages over direct tumor tissue biopsy, with faster, less invasive sample collection^5^. Identified variants are likely to present a better representation across heterogeneous tumors^6^. Additionally, for MRD monitoring and early cancer detection, where tumor tissue may be unavailable, liquid biopsy becomes the preferred choice.

The decreasing cost and increasing throughput of next-generation sequencing technology have facilitated the development of assays directly assessing ctDNA. The high throughput of existing short-read sequencing platforms is particularly suitable for high-depth multi-gene panels aiming to detect variants at low allelic fractions (AFs). Consequently, NGS ctDNA assays are rapidly gaining adoption in precision oncology applications^7,8^. Challenges persist, mainly due to limited sample volumes and the number of DNA fragments required for reliable detection of ultra-low AF variants. For instance, a single tube of blood typically yields less than 4mL of plasma, from which less than 30ng of DNA (∼9,000 genome copies) can be extracted. Material loss during library construction, target capture, and sequencing further limits the sequenced genome copies to fewer than 6000, even at saturation sequencing depths.

While the overall base accuracy is commendable, the prevalent short-read sequencing platform exhibits an error rate of as high as 1 in 1000 bases, akin to the allele frequency of targeted somatic variants. To enhance accuracy and discernment between background errors and real somatic variants, some ctDNA assays have incorporated Unique Molecular Identifiers (UMIs) in the pipeline^9^. UMIs introduce distinctive tags (base sequences) at the ends of template DNA molecules before PCR amplification. These uniquely tagged templates undergo amplification and sequencing multiple times and enable grouping reads based on their original molecules.

However, some assays opt not to include UMIs or reach higher sequencing depth, primarily due to cost considerations, especially for late-stage treatment selection where the sample tumor fraction is relatively high. In such cases, where error correction and high accuracy still remain imperative, a non-UMI-based consensus deduplication method can be instrumental. Specifically, reads with the same start and end mapped position can be intelligently identified and computationally collapsed, significantly mitigating the impact of PCR duplicates and sequencing errors in the variant calling process.

A typical ctDNA NGS analysis pipeline encompasses three crucial steps: alignment, consensus generation, and variant calling. While BWA-mem is widely adopted as the mainstream aligner for short reads, a consensus on best practices has yet to be established for the other two steps. Several pipelines integrate consensus generation and variant calling within a single module, exemplified by tools like “DeepSNVMiner”^10^, “MAGERI”^11^, “smCounter”^12^, and “smCounter2”^13^. On the other hand, some pipelines opt for separate consensus generation and variant calling tools, offering more flexibility and easier adaptation to new data types. Popular consensus generation tools include “Fgbio”^14^, “Picard”^15^, “Gencore”^16^, and Sentieon’s UMI analyzing pipeline^17^. Fgbio is typically applied to datasets with UMIs, Picard serves as a traditional deduplication tool, selecting reads with the best quality score as group representation. Gencore and Sentieon UMI pipeline, on the other hand, is described as a consensus-based deduplication tool that works with or without UMIs, providing higher accuracy compared to alternatives. For somatic variant calling, popular tools include “Mutect2”^18^, “Vardict”^19^, “Strelka2”^20^, and “VarScan2”^21^. While these callers were not originally designed for ctDNA assays, a recent benchmark study^13,22^ demonstrated their reasonable accuracy in this context.

The Genome in a Bottle (GIAB) consortium, led by NIST, has made significant strides in providing reliable reference materials and benchmark datasets for germline variant calling, with the first of these datasets becoming available in 2014^23^. However, progress in somatic variant benchmarking has been slower, primarily due to the absence of publicly available reference materials and benchmark datasets until very recently. The Sequencing Quality Control2 (SEQC2) project, published in 2021, represents a pioneering multi-center effort that offers a comprehensive reference dataset and truth for tumor tissue^24,25^ and ctDNA samples^22^. Among the ctDNA samples, some of the generated datasets contain variants with a median Allele Frequency (AF) at 0.5-1%. Their AFs fall within the scope of treatment selection assays, making them particularly relevant for the purposes of this research. Therefore, these datasets and are chosen and as a benchmark dataset for this study.

This study introduces Sentieon’s non-UMI ctDNA pipeline, offering a rapid and accurate solution for small somatic variant calling from ctDNA NGS data. The pipeline comprises four core modules: 1) Sentieon BWA-turbo: This module, a machine learning-guided accelerated version of BWA-mem for alignment^26^. 2) Sentieon Consensus-based Deduplication module: This tool facilitates consensus generation by grouping and generating consensus reads based on read position^27^. 3) TNscope: A haplotype-based somatic variant caller that adheres to the principles of mathematical models first implemented in the GATK HaplotypeCaller and Mutect2. It demonstrates higher sensitivity to low Allele Frequency (AF) variants and incorporates various other improvements^28^. 4) TNscope Filter: This module serves as a customizable filtering tool designed to remove false-positive variants.

## Result

In this study, we assess the accuracy of the Sentieon non-UMI ctDNA pipeline using three distinct datasets: two in-vitro mixtures with known ground truth and one clinical dataset derived from real-world diagnostic samples. To gauge the accuracy of the Sentieon ctDNA pipeline, we employ two in-vitro mixed datasets with established truthset variant calls. We calculate variant calling accuracy using the known truth and compare Sentieon’s accuracy to alternative variant callers.

Furthermore, we apply the Sentieon pipeline to a real clinical dataset obtained from samples of lung cancer and colorectal cancer. This evaluation aims to understand the performance of the Sentieon pipeline in a practical diagnostic setting.

### SEQC2 ctDNA Dataset

Firstly, we incorporated data from the recently published SEQC2 dataset^21^ to benchmark the Sentieon non-UMI ctDNA pipeline. The SEQC2 study, a large multi-center project, aims to generate ctDNA reference samples and benchmark currently available ctDNA assays. As part of this effort, the SEQC2 project created two reference samples through in-vitro mixtures at different ratios of two samples with known somatic variants.

The Lbx-high mixture encompasses variants with a median frequency of approximately 1%, with a majority exceeding 0.5%. Conversely, the Lbx-low mixture contains variants with a median frequency of around 0.2%, with a majority surpassing 0.1%. Both reference DNAs underwent sequencing and bioinformatics analysis by multiple ctDNA assay vendors. Notably, the BRP (Burning Rock Biotech) assay claimed to have the highest accuracy in the project, leading us to select the BRP datasets for our benchmark. Detailed information about the dataset preparation methods can be found in the SEQC2 study paper^21^ (Fig 2).

**Figure 1.**
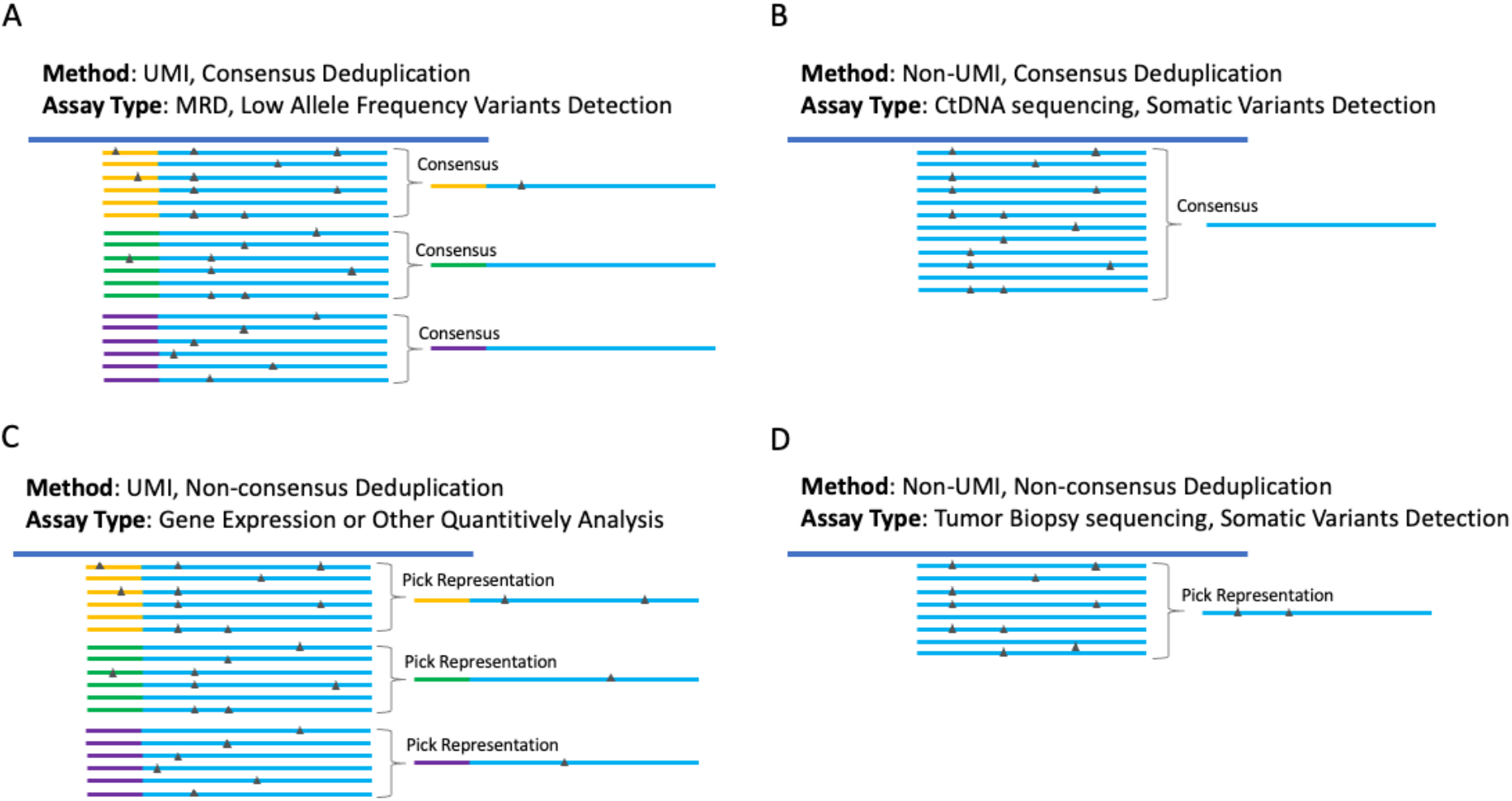
Deduplication in the analysis pipeline serves to eliminate errors introduced by PCR or sequencing, as well as manage PCR duplicates. Whether adding UMI or conducting consensus-based deduplication should be determined by dataset type and assay purpose. Four example scenarios are showed in the figure, with blue line representing sequenced reads, colorful line representing UMI tags, black triangles representing errors.

**Figure 2.**
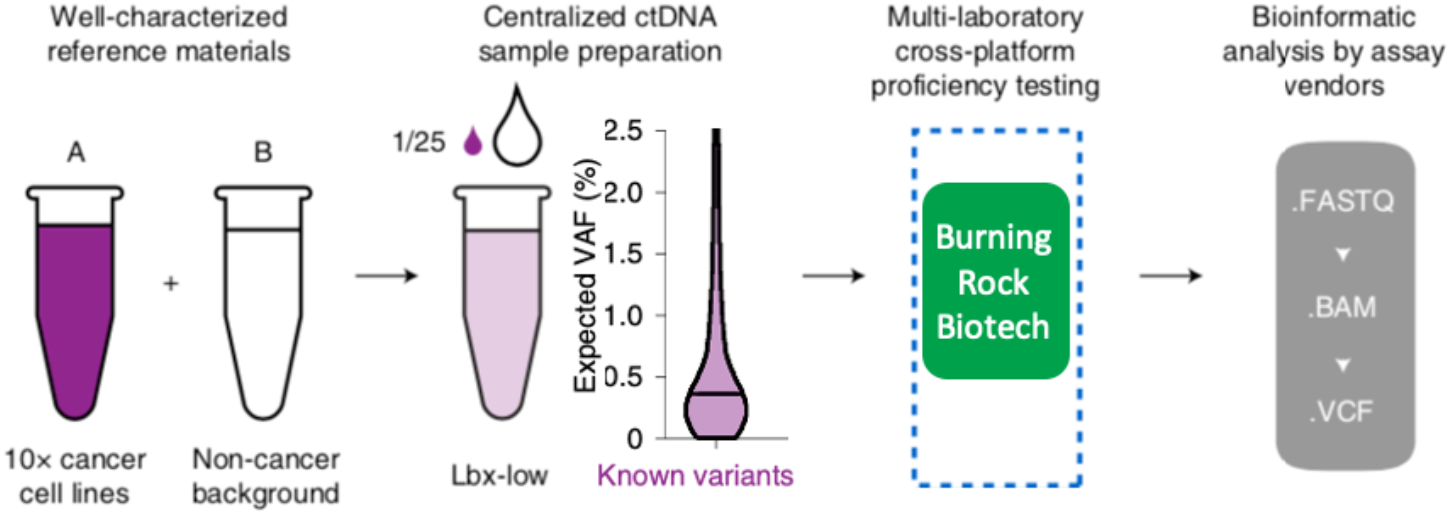
Schematic Diagram illustrating the reference sample generation used in this section, assay process, expected variant AFs, and variant calling. Modified from publication^21^.

For this benchmark project, we opted to focus on the eight Lbx-high samples, as their variants have AFs within a range detectable without the need for UMI. The fastq files for these samples were subsampled from their original depth to a level lower than 10,000x, and analysis was conducted without considering UMIs information. This subsampled depth is likely more reflective of real-world cost constraints for production assays.

Then the treated BAM files were processed by Sentieon pipeline for consensus-based deduplication and variant calling, and the accuracy of variant calling was assessed using the SEQC2 truthset. Meanwhile, the original depth with UMI datasets were processed by BRP’s own pipeline which relies on a UMI consensus tool developed by BRP and VarScan2^21^ (supp note). Then accuracy values of both pipelines were calculated and compared head-to-head.

Figure above illustrates the Precision and Recall for the two pipelines. Surprisingly, recall and precision is not affected by the lower sequencing depth and lack of UMI info, after being processed by Sentieon pipeline. The disparity in the overall F1 score is not substantial, suggesting that it might be adequate for treatment selection purposes as a more cost-effective assay.

### In-Vitro Mixture of DNA from Healthy Individuals

The next benchmark dataset originates from a blend of genomic DNA (gDNA) or cell-free DNA (cfDNA) extracted from two healthy individuals. For each of these individuals, truthset variants were determined using the GATK germline variant calling pipeline. Subsequently, DNA from one individual (spike-in) was mixed at a 1% titration rate with DNA from another individual (background) (Fig 4). Libraries were constructed from these mixed DNA samples with a 10ng input. A customized panel probe hybridization was applied to all libraries, resulting in 57 homozygous SNPs at a 1% AF and 70 heterozygous truth SNPs at a 0.5% AF within the panel region.

**Figure 3.**
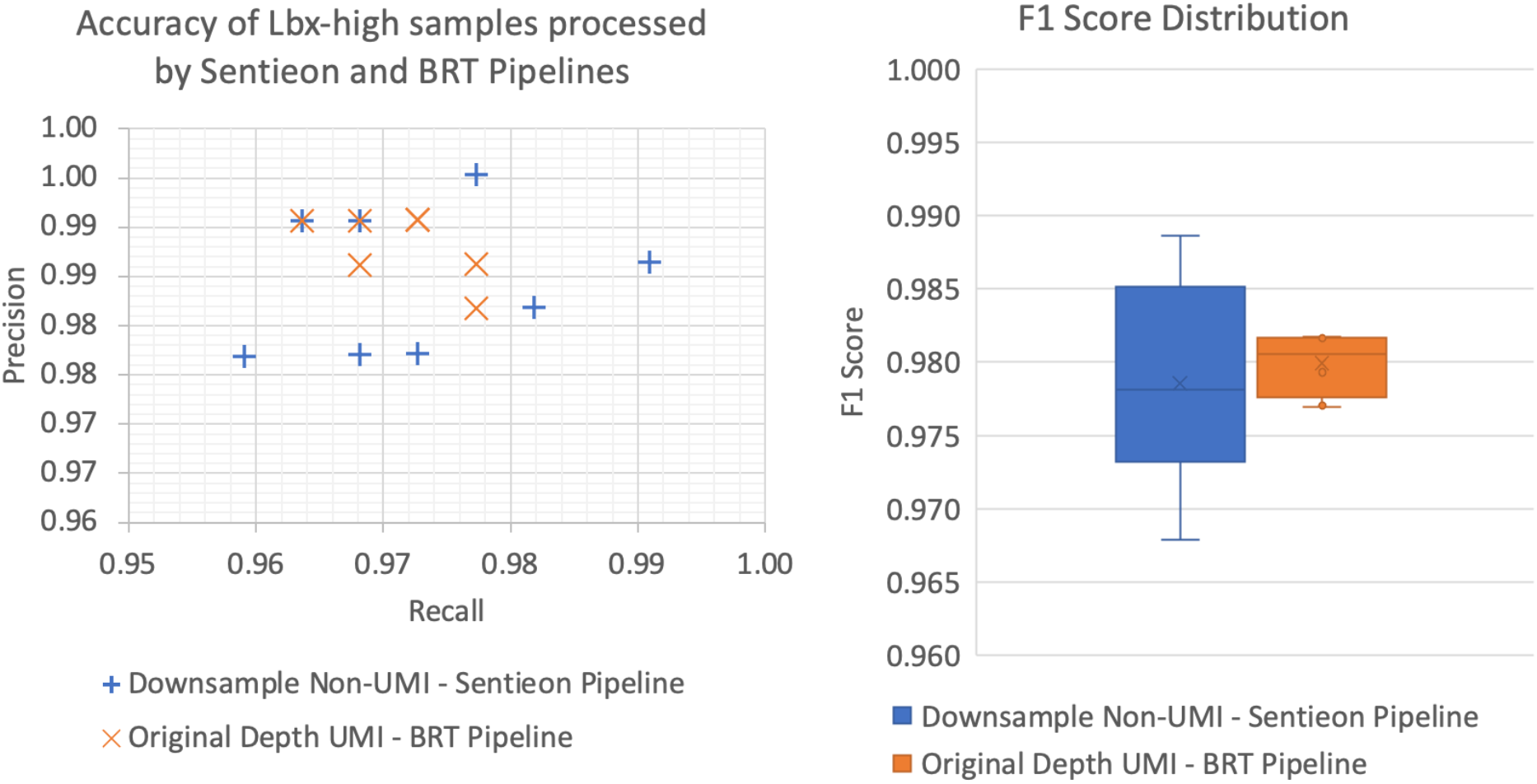
Precisions vs. Recalls of Lbx-high samples processed by Sentieon and alternative pipeline; F1 score distribution of two pipelines.

**Figure 4.**
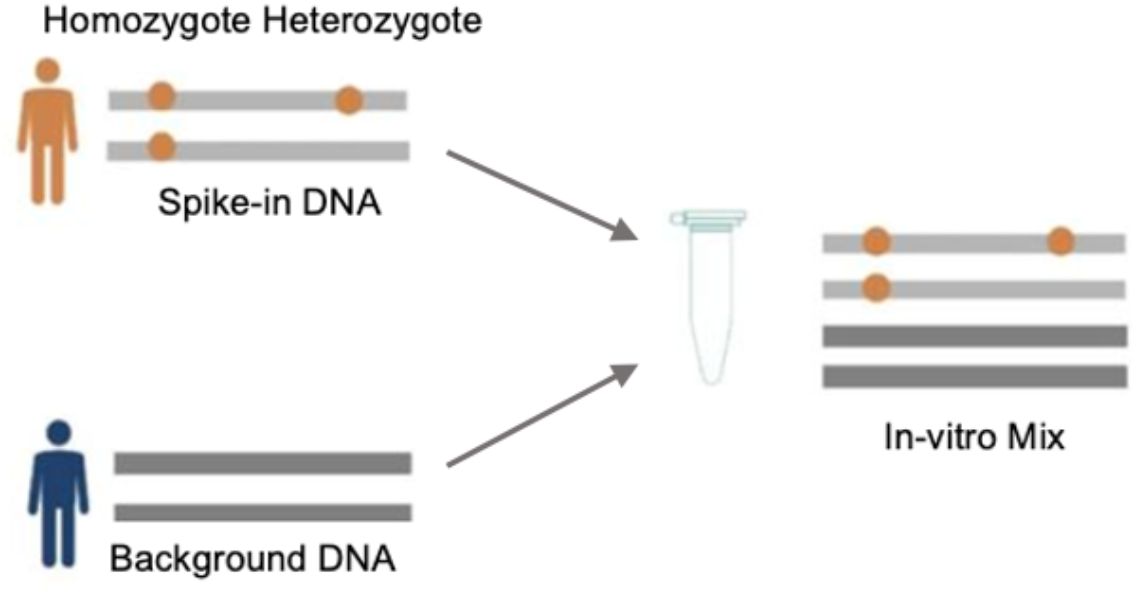
In-Vitro mixture samples were prepared by titration of DNAs from two healthy individuals.

Following this, the libraries were sequenced on Illumina or MGI platforms, and the datasets were subsampled to 8,000x before analysis. Due to data availability, two of the titration datasets (Lib74 and NL190929-1C) were examined, and detailed information for each dataset is provided in Table 2, with the detailed wet lab and data processing described in the methods section.

**Table 1.** Pre-dedup and post-dedup depth of the evaluated datasets.

| Run | ST | Lib | Pre-Dedup Depth | Subsampled Pre-Dedup Depth | Subsampled Post-Dedup Depth |
| --- | --- | --- | --- | --- | --- |
| SRR13200966 | 25 | 1 | 42,995x | 6,877x | 3,293x |
| SRR13200965 | 25 | 2 | 36,078x | 5,772x | 3,048x |
| SRR13201010 | 25 | 3 | 42,187x | 6,747x | 3,276x |
| SRR13201009 | 25 | 4 | 49,339x | 7,894x | 3,259x |
| SRR13201008 | 26 | 1 | 49,776x | 7,966x | 3,660x |
| SRR13201007 | 26 | 2 | 65,458x | 10,474x | 4,126x |
| SRR13201006 | 26 | 3 | 51,610x | 8,257x | 3,683x |
| SRR13201005 | 26 | 4 | 50,346x | 8,054x | 3,705x |

**Table 2.**
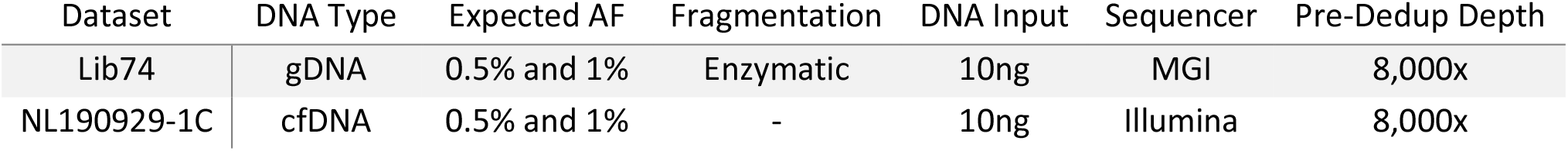
Sample, library, and sequencing settings of the in-vitro mixture datasets.

The 1% titration datasets were designed to mimic biopsy and sequencing for treatment selection purpose assay. UMIs in these datasets were initially included but deliberately ignored during analysis to align more closely with real-world applications. To assess the performance of the analysis pipeline across diverse wet lab settings, DNA types, and sequencing platforms, we tried to employ varied combinations. Both datasets underwent processing by the Sentieon ctDNA pipeline, and we also applied an alternative pipeline using “Fgbio + Vardict” to generate baseline performance. The resulting variant calls were then compared against the truthset to calculate accuracy.

Although both the Sentieon and alternative pipelines demonstrated high accuracy (Fig 5), the Sentieon pipeline exhibited higher recalls, particularly at 0.5% AF variants, effectively enabling a lower limit of detection. While precision values were comparable for both pipelines, Sentieon’s overall F1 Score consistently outperformed that of the alternative pipeline.

**Figure 5.**
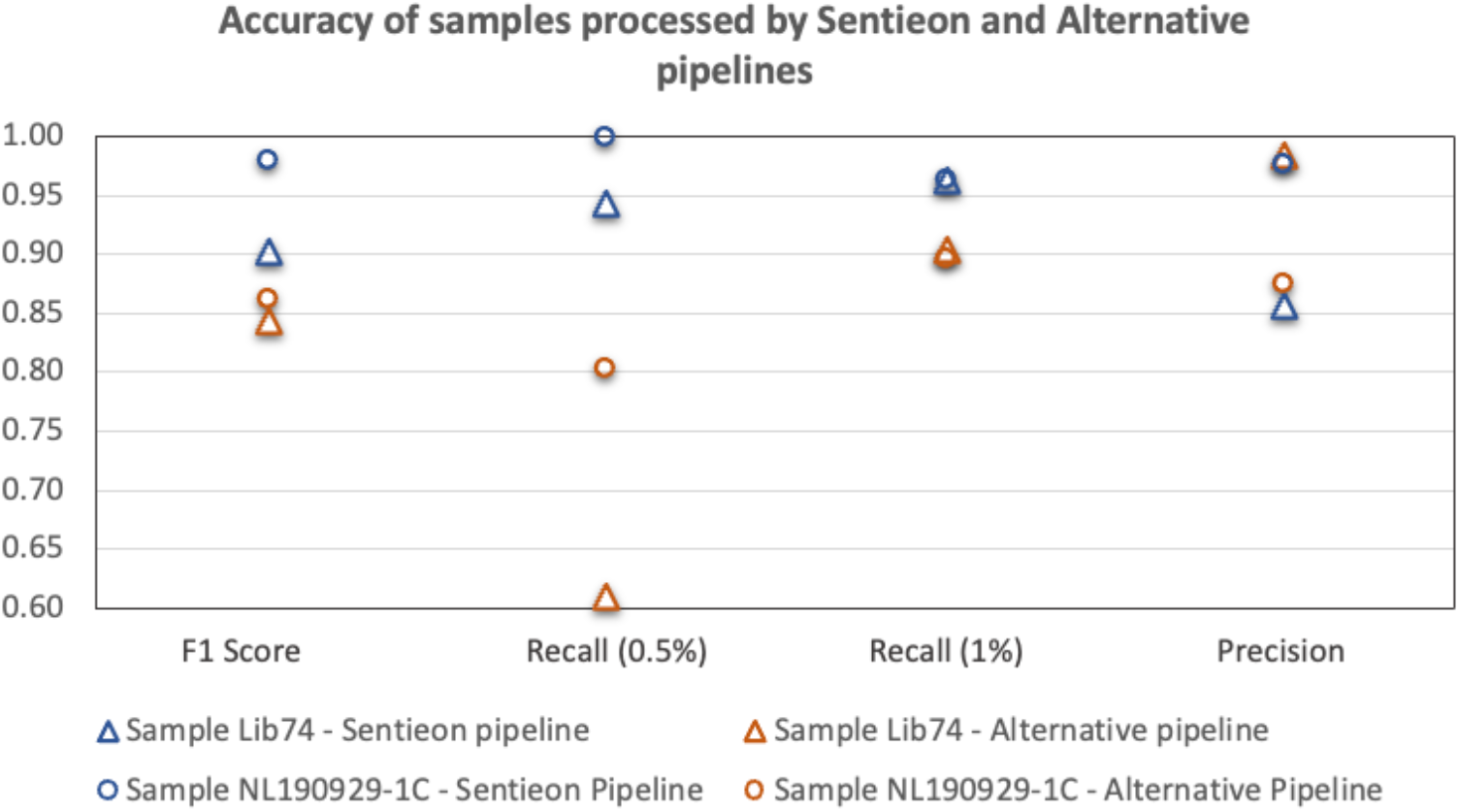
Evaluating the accuracy of the Sentieon ctDNA pipeline and an alternative pipeline on in-vitro mixture datasets. Recall was calculated separately for 0.5% AF variants and 1% AF variants.

**Figure 6.**
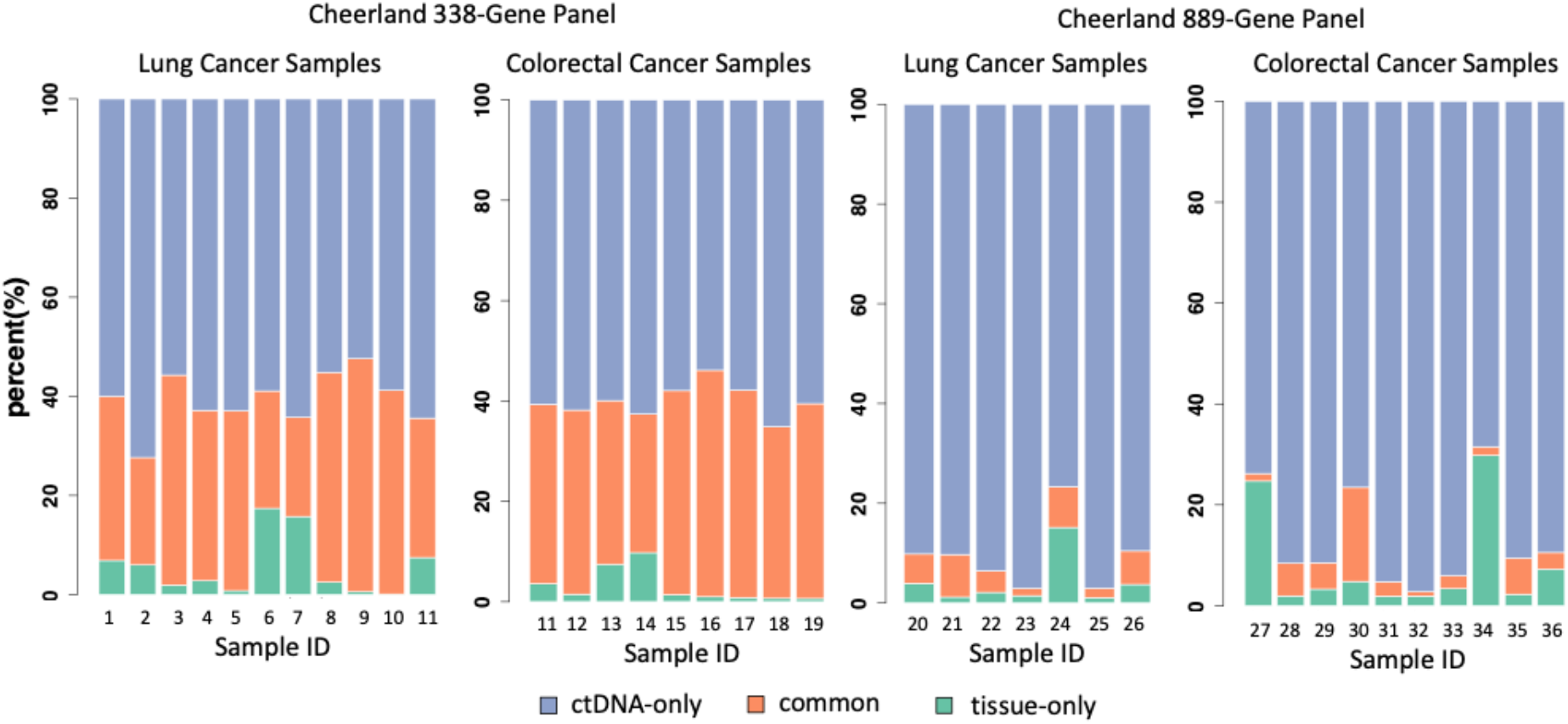
Comparing ctDNA variant detection in ctDNA (>0.5% AF) samples and tumor tissue samples (>5% AF) from the same patient. Cancer type and panel type are indicated above each bar chart. Different colors represent ctDNA-only, tissue-only, and common variants.

**Figure 7.**
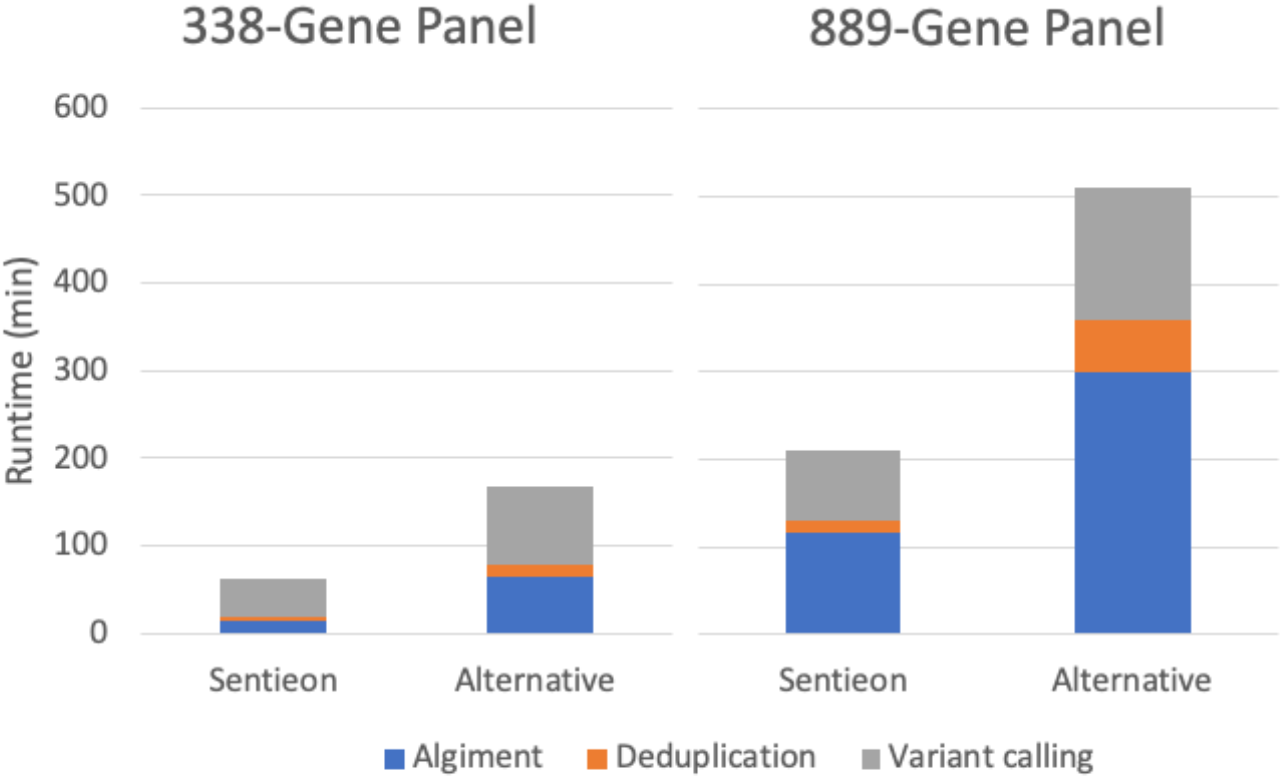
FASTQ to VCF Runtime benchmark. Computation was done using 6 threads and less than 32GB RAM. Runtime was generated from averaging processing time of 5 datasets.

### Clinical Cancer Sample Validation

The previous analyses highlight the robust performance of the Sentieon ctDNA pipeline on in-vitro mixtures datasets. To further evaluate the pipeline’s capabilities, we conducted tests on real-world clinical samples, comprising a total of 36 samples from lung cancer and colorectal cancer patients. These samples were processed by Cheerland Biotechnology, using the Cheerland 338-gene personalized treatment selection and the 889-gene comprehensive panel. The Sentieon non-UMI ctDNA pipeline was employed for somatic variant calling.

Each sample had a pre-determined set of oncogenic clonal mutations with AF higher than 5%, identified through tumor tissue biopsy sequencing for the same patient. Subsequently, variants identified in the ctDNA assays (>0.5% AF) were cross-compared with the tumor biopsy variant set to distinguish whether a variant is ctDNA-only, tissue-only, or present in both samples.

It’s important to note that, due to the large panel coverage, ctDNA-only variants may include non-tumor somatic variants (e.g., clonal hematopoiesis), or subclone variants omitted from heterogeneous tumor tissue samples. Conversely, tissue-only variants may indicate the growth location of the biopsied subclone tissue that does not have access to the main blood vessel, shedding a lower amount of ctDNA into the circulation system.

While determining the true positive rate for clinical samples is challenging, the cross-comparison of tissue and ctDNA samples from the same patient provides valuable evidence of the assay’s accuracy. Notably, lung cancer and colorectal cancer samples exhibited no significant difference in the percentage of ctDNA-only variants, common variants, and low-percentage tissue samples. This suggests that the 338-gene panel design and Sentieon pipeline are adaptable to both lung and colorectal cancer samples, and potentially other cancer types as well.

In contrast, the 889-gene panel showed a significantly higher percentage of ctDNA-only variants. This may be attributed to the wider gene coverage detecting variants not included in the heterogeneous tissue sample or even non-tumor somatic variants. The results indicate that the 338-gene panel is more focused on detecting variants originally present in tumor tissue, while the 889-gene panel provides a greater number of variants and comprehensive information on patients to identify and manage all potential health risks.

Cheerland Biotechnology offers an extensive range of nearly 50 diagnostic assays, catering to various crucial tumor types such as lung cancer, colorectal cancer, breast cancer, gynecologic tumors, liver and bile tumors, digestive system tumors, urinary system tumors, thyroid cancer, brain glioma, and more. The capture-based assay comprises a small panel (338 genes) and a large panel (889 genes).

To date, over 80,000 samples have been received and successfully processed. In 2023 alone, this included approximately 14,000 cases using the small panel and 6,000 cases utilizing the large panel. In the preceding year, 2022, 11,000 small panel cases and 6,000 large panel cases were processed. All datasets generated from these assays have been meticulously stored and are actively contributing to clinical science and biomedical translational research.

### Runtime Comparison

The Sentieon non-UMI ctDNA pipeline is fine-tuned for processing a substantial volume of reads while upholding high accuracy for consensus reads. Its efficient algorithm and implementation in C/C++ result in a processing speed approximately 4 times faster than Picard in the speed benchmark. The overall pipeline from FASTQ to VCF demonstrated a notable acceleration of 2-3 times. This benchmark involved the use of 10 datasets from Cheerland 338-gene panel and 889-gene panel datasets, with the allocation of 6 threads and 32GB RAM per run on a logical core Intel Xeon platform.

## Methods

### Generation of Healthy Individuals Invitro-mix Dataset

Genomic DNAs were isolated from blood samples of two healthy individuals using the TIANamp Genomic DNA kit (#DP304, TIANGEN BIOTECH). Simultaneously, cell-free DNAs were extracted from the same individuals employing the Serum/Plasma Circulating DNA Kit (#DP339, TIANGEN BIOTECH).

Genomic DNA or cfDNA from one individual (spike-in) was blended with DNA from another individual (background) at a 1% titration rate. Library 74 was crafted using the NadPrep DNA Library Preparation Kit (for MGI), incorporating the Bi-Molecular Identifier (BMI) adapter (#1003821, Nanodigmbio), from 10ng gDNA fragmented by NEBNext dsDNA Fragmentase (#M0348, NEB). Library NL190929-1c was generated with the NadPrep DNA Library Preparation Kit (for Illumina), including the UMI adapter (#1003431, Nanodigmbio), from 10ng cfDNA. Following end repair and A-tailing, dilution was performed. A 40μL volume of NadPrep SP Beads was employed for library cleanup, and ligated fragments were amplified with 6-9 cycles using a 0.5M index primers mix. Library yields were maintained between 500-1000ng. A 60μL volume of NadPrep SP Beads was used to recycle libraries.

Both libraries underwent processing through single-tube capture hybridization. Following the Nanodigmbio hybridization capture protocol (NadPrep Hybrid Capture Reagents, #1005101), each pool of DNA was combined with 5μL of 1mg Cot-1 DNA (Invitrogen) and 2μL of 0.2nmol NadPrep NanoBlockers (#1006204 for MGI, #1006101 for Illumina, Nanodigmbio) to prevent cross-hybridization and minimize off-target capture. The library and blocker were dried and re-suspended in hybridization buffer and enhancer. Target capture with 38kb “SNP ID” Panel Probes (Nanodigmbio) was carried out overnight. Streptavidin M270 (Invitrogen) beads were utilized to isolate hybridized targets following the Nanodigmbio hybridization capture protocol. Captured DNA fragments were then amplified with 13–15 cycles of PCR.

Subsequently, the libraries were sequenced using either 100bp paired-end runs on the MGI-2000 sequencer at the Nanodigmbio R&D Center or 150bp paired-end runs on the Illumina XTen sequencer at the Wuxi Genome Center. Germline variants of the two individuals were identified using the GATK pipeline, and germline variants unique to two contributor samples served as the truthset for accuracy calculation.

### SEQC2 Variants Accuracy Benchmark

VCF files generated by either the Sentieon pipeline or BRP pipeline underwent filtering using both the SEQC2 truth bed file and BRP panel bed file. Specifically, the intersection of CTR_hg19.bed and BRP2.bed produced the CTR_hg19_BRP2.bed interval. Subsequently, the PASS VCF was queried against the truth VCF within the CTR_hg19_BRP2.bed regions to determine the counts of True Positives and False Negatives.

Similarly, the intersection of KnownNegatives_hg19.bed and BRP2.bed resulted in the KnownNegatives_hg19_BRP2.bed interval. PASS variants located within the KnownNegatives_hg19_BRP2.bed regions were then counted to determine the number of False Positives. The comparison of VCF files was facilitated using RTG Tools (v3.8.2).

### Sentieon ctDNA Pipeline - Alignment and Sort

The Sentieon Genomics tool set constitutes a suite of software tools designed for the secondary analysis of next-generation sequence data. The Sentieon pipelines feature optimized implementations of the mathematical models employed in highly accurate variant calling pipelines. Performance enhancements are achieved through algorithm optimization and improved compute resource management.

Prior to variant calling, FASTQ data undergo alignment to the human reference genome using Sentieon BWA-turbo. BWA-turbo is developed by Sentieon as a machine learning-guided and optimized version of BWA-mem, resulting in an approximate 4-6x speedup while maintaining identical alignments for all high-quality reads. BWA-turbo strategically allocates computational resources to reads likely to contribute to downstream variant calling, thereby improving computational efficiency without negatively impacting variant calling accuracy. Read coordinate sorting was carried out using the Sentieon utility module “sort.”

### Consensus Generation

Generating a consensus read in the deduplication step is considered a crucial analysis stage for high-depth liquid biopsy sequencing. Downstream variant calling relies on deduplication to prevent double counting of evidence. In addition to accurately generating the consensus read base sequence, correctly modeling the confidence in the base, and outputting this information in the consensus base quality are essential for downstream analysis as well. The base quality assigned to the consensus bases should precisely reflect the underlying probability of a base error.

The Sentieon Deduplication module function in non-consensus mode (similar to Picard) or consensus mode, with or without UMI^27^. The consensus module employs sophisticated modeling of base errors introduced during library construction and the sequencing process to enhance consensus read accuracy. Reads are grouped based on their positions and UMI tags if available, and the grouped reads are statistically modeled to account for multiple sources of base error. Parameters of the statistical model are learned and calibrated directly from the dataset without user input. Consensus reads are then called from the grouped reads using a Bayesian model informed by the learned parameters, and overlapping read pairs are merged. It’s important to note that during consensus calling, no reads are discarded, and all read information is considered for model calibration. The confidence in consensus calling is reflected in the assigned base quality of consensus bases, enhancing downstream callers’ ability to improve accuracy.

### TNscope and TNscope-filter

TNscope is a haplotype-based variant caller that adheres to the fundamental principles of mathematical models initially implemented in the GATK HaplotypeCaller and MuTect2. This approach involves active region detection, assembly of haplotypes from the reference and local read data using a de Bruijn-like graph, and the use of a pair-HMM for calculating read-haplotype likelihoods followed by genotype assignment. Similar to MuTect2, TNscope evaluates tumor and normal haplotypes jointly when a matching normal sample is available, leading to significantly higher precision in somatic variant detection.

Several enhancements have been integrated into the mathematical model of TNscope to improve both its recall and precision. The highly efficient implementation allows TNscope to choose a lower threshold for triggering active regions, enabling a more comprehensive evaluation of potential variants. Additionally, the detected active regions are typically of higher quality as TNscope employs a statistical model to trigger active regions rather than a fixed cutoff. Local assembly has been improved too, resulting in more frequent identification of the correct variant haplotype. Genotyping has also been enhanced through the adoption of a novel quality score model and various nonparametric statistical tests to eliminate false-positive variants. For high-depth targeted sequencing data, down sampling is not required due to TNscope’s computational efficiency, making it an ideal haplotype-based variant caller for detecting rare somatic events in ctDNA samples.

TNscope provides several novel variant annotations that contribute to improved variant filtration. The TNscope-filter module serves as a VCF filtering tool, fully utilizing these annotations and identifying false-positive variants from raw VCF files based on input parameters. Its functionality is akin to BCFtools^29^ but is more compatible with other Sentieon modules.

## Conclusions

In this study, we introduce the Sentieon non-UMI ctDNA analysis pipeline and assess its accuracy using both in-vitro mixture and real clinical datasets. Starting from the same benchmark datasets, we observed superior recall and precision from Sentieon pipeline compared to alternative pipelines, or comparable accuracy from a dataset generated with lower cost. The exceptional performance of the Sentieon pipeline can be attributed to the sophisticated statistical model employed in the consensus generation tool, coupled with highly accurate somatic variant calling from Sentieon TNscope. In addition to its accuracy, the Sentieon ctDNA pipeline also demonstrates significantly faster processing times than alternative pipelines, facilitating the timely analysis of high-depth large-panel datasets.

## Code Availability

Script of Sentieon non-UMI pipelines benchmarked in this study can be found on Github page: https://github.com/Sentieon/sentieon-scripts/blob/master/example_pipelines/somatic/TNscope/Somatic_ctDNA_without_UMI.sh

